# A miRNA-mediated gene regulatory network supports seasonal plasticity in the temperate coral *Astrangia poculata*

**DOI:** 10.64898/2026.06.11.731608

**Authors:** Jill Ashey, Chloé Gilligan, Hollie M. Putnam

**Affiliations:** Department of Biological Sciences, University of Rhode Island, Kingston RI USA; Department of Biology, University of Pennsylvania, Philadelphia PA USA; Department of Biology, University of Massachusetts, Dartmouth MA USA

## Abstract

Phenotypic plasticity is a critical strategy for sessile marine invertebrates that cannot escape changing environmental conditions. For corals facing intensifying climate change, the molecular mechanisms that generate and regulate acclimatory responses are central to understanding performance and persistence. MicroRNAs (miRNAs), small non-coding RNAs that regulate gene expression via translational repression and transcript degradation, are compelling candidates for mediating such plasticity, but their role in coral biology remains poorly characterized. Here, we use *Astrangia poculata*, a temperate coral that endures annual temperature ranges exceeding 25°C, to investigate the potential for miRNA-mediated plasticity across seasonal and thermal contexts. We exposed adult aposymbiotic colonies to ambient seasonal temperatures (∼5–22°C) and a chronic +3°C warming treatment from February to August 2021, sampling monthly for physiological analyses and at three time points (February, June, August) for molecular analyses. Seasonal change drove significant shifts in photosynthesis, respiration, and soluble protein, whereas the +3°C treatment had minimal physiological effect. RNA-seq analysis identified the strongest response across seasons, with transcriptional functional enrichment shifting from protein homeostasis, and cellular integrity in the winter, to immunity, metabolism, and reproduction in the summer. We identified 51 miRNAs in *A. poculata*, 46 of which are novel to this species, providing the first characterization of the miRNA repertoire in this species. Target prediction and co-expression analyses revealed that while mRNA-miRNA networks maintain a stable infrastructure across seasons and treatments, specific interactions are rewired to drive seasonal biology. This dynamic regulation suppresses energy-intensive cell division and morphogenesis during winter quiescence, while shifting to regulate tissue remodeling and reproduction genes during the summer. These results establish miRNAs as seasonal gene regulators in a temperate coral and suggest that the molecular infrastructure underlying its plasticity may also confer resilience to moderate thermal stress.

## Introduction

Regulatory physiology enables organisms to make dynamic adjustments in response to changing environments, maintaining stable internal conditions optimized for cellular function (Bernard, 1865; Cannon, 1929). This understanding of the “milieu intérieur” (Bernard, 1865) has developed into the modern study of plasticity and its underlying regulatory mechanisms (Hofmann & Todgham, 2010; Wingfield, 2013). Phenotypic plasticity, the ability of an organism to adjust its phenotype when exposed to variation in the environment, is a common strategy for coping with such variation (West-Eberhard, 1989). This plasticity is often most pronounced under predictable seasonal cycles, illustrated by classic examples such as coat color shifts in snowshoe hairs (Zimova et al., 2018), organ remodeling in migratory birds (Piersma & Van Gils, 2011), and metabolic suppression during torpor and hibernation (Van Breukelen & Martin, 2015). While these macro-level physiological shifts are well documented, the molecular foundations that coordinate these seasonal responses remain less explored.

Phenotypic plasticity can be driven by changes in the transcriptome, where messenger RNA (mRNA) can be rapidly synthesized or regulated to enable the translation of proteins for dynamic responses to environmental signals. While many studies have explored transcriptomic responses to high frequency environmental fluctuations, seasonal change, and acute and chronic stressors (Gracey, 2007; Hofmann & Todgham, 2010; Evans & Hofmann, 2012), the mechanisms that regulate these gene activity states remain less understood, particularly in non-model marine taxa (Eirin-Lopez & Putnam, 2019; Johnson & Wong, 2026). This regulation often occurs through epigenetic mechanisms, which affect changes in gene expression without changes to the underlying DNA sequence (Bird, 2007; Costa, 2008; Feil & Fraga, 2012). One such mechanism is microRNAs (miRNAs), small non-coding RNAs that typically bind to target mRNAs to facilitate translational repression or transcript degradation (O’Brien et al., 2018), making them critical mediators of gene expression changes and thus plasticity.

For sessile marine invertebrates, which are literally anchored to their environment and cannot move to avoid adverse conditions, phenotypic and physiological flexibility is a critical mechanism to avoid mortality (Sanford & Kelly, 2011; Padilla & Savedo, 2013). Thus, regulation at the cellular and molecular levels is essential to life in a changing climate for these sessile taxa. Scleractinian corals represent a compelling model for studying phenotypic plasticity (Todd, 2008) and its regulation under climate change (Putnam, 2021). Not only must corals maintain physiological homeostasis under naturally fluctuating temperature, light, and resource availability (Brown, 1997), but they are also now additionally challenged by anthropogenic climate change (Hughes et al., 2017).

A major acclimatory response of corals in the Anthropocene is dealing with the cellular consequences of increasing temperature exposure (Helgoe et al., 2024). The coral-algal nutritional symbiosis is based on carbon and nitrogen exchanges between the partners (Muscatine & Porter, 1977), where the symbionts photosynthesize and generate >90% of the daily metabolic demand of the host (Falkowski et al., 1984; Muscatine et al., 1981), and the host provides nitrogen from heterotrophy for algal protein turnover and population growth (Falkowski et al., 1993). Thus, as increasing temperatures break down this symbiosis, corals must manage challenges of fluctuating carbon and nitrogen levels (Rädecker et al., 2021), acid-base imbalances (Tresguerres et al., 2017), and reactive oxygen species damage (Weis, 2008), among other homeostatic disruptions (Helgoe et al., 2024). In light of this, many studies have explored coral transcriptomic responses (reviewed in Drury, 2020) to high frequency temperature fluctuations, seasonal thermal change, and acute and chronic temperature stress. In contrast, the exploration of miRNAs regulation of gene expression in corals remains limited. Initial evidence suggests that miRNAs have roles in core coral biological aspects (Ashey, Rodriguez-Casariego, et al., 2025), including calcification (Liew et al., 2014), stress responses (Despard et al., 2025; Gajigan & Conaco, 2017), and symbiosis (Baumgarten et al., 2018). Given the emerging evidence for miRNAs to regulate gene expression in multiple coral taxa (Ashey et al., 2025), juxtaposed with the growing environmental stress of climate change, regulatory physiology is essential to our understanding of coral performance under current and future environmental conditions.

Temperate coral species–such as the Northern star coral, *Astrangia poculata*–exhibit extreme physiological flexibility across seasons (reviewed in Ashey et al., 2025). As a facultatively symbiotic coral, *A. poculata* exists across a continuum of symbiotic states, ranging from brown pigmented colonies with high symbiont densities to nearly aposymbiotic (white) individuals (Ashey et al., 2025). Northern populations of *A. poculata* endure annual temperature changes exceeding 25°C, during which they transition from a state of winter quiescence (metabolic dormancy; Grace, 2017) to an active summer period of growth and reproduction (Dimond & Carrington, 2007; Szmant-Froelich et al., 1980) at 20–25°C. Collectively, these coral and holobiont responses of *A*. *poculata* under extreme and recurrent seasonal changes indicate its capacity to effectively shift its physiology to match environmental conditions (Dimond & Carrington, 2007, 2008; Grace, 2017; Sharp et al., 2017; Trumbauer et al., 2021, 2022) and suggest regulatory mechanisms capable of coordinating these shifts.

Narragansett Bay, Rhode Island, home to a high density of *A*. *poculata* colonies (Lindsay et al., 2025), is warming at a rapid rate, with long-term environmental data showing significant increases in seawater temperature since 1950 (Collie et al., 2008; Fulweiler et al., 2015; Humphries et al., 2022). Further temperature increase is predicted under climate change, leading to higher average seawater temperatures year-round and potentially altering seasonal temperature ranges that *A. poculata* has historically experienced (Fulweiler et al., 2015; Humphries et al., 2022; Smith et al., 2010), which could exceed existing acclimatory mechanisms. Here, we use the *A. poculata* as a model for acclimatory phenotypic plasticity and regulatory physiology to test potential for miRNA-mediated regulation of gene expression and plasticity under seasonal and elevated thermal regimes.

We exposed adult *A. poculata* colonies to ambient seasonal temperatures (5–22°C) as well as a chronic thermal increase treatment (+3°C above ambient) from February to August, which spans the lower and upper temperatures of their thermal range. Physiological responses were measured monthly, and molecular samples were collected at three time points. We hypothesized that *A. poculata* would display increased metabolic demands, and reduced carbohydrate and protein content in the warmest month (August) compared to the coldest month (February), and that this transition in metabolism and energy stores would be linked to changes in gene expression, regulated by miRNAs. Further, we hypothesized that these seasonal patterns would be intensified by the +3°C temperature increase, driving higher gene expression regulation as temperatures reached their seasonal peak.

## Methods

### Collection, Acclimation, and Experimental Design

This experiment was conducted from December 2020 to August 2021, with the experimental exposure period being from February 2021 to August 2021. In December 2020, adult white phenotype *A. poculata* were collected from Fort Wetherill (41.477371, −71.360113) and the Graduate School of Oceanography (GSO) dock (41.49231,-71.41883) in Narragansett Bay, Narragansett, RI, USA. Following collection, corals were transported to the Graduate School of Oceanography (GSO) at the University of Rhode Island and placed in glass, flow-through experimental tanks (35 L; Aqueon Products) for a two month acclimation period under ambient conditions. Seawater was pumped from Narragansett Bay and passed through a sand filter (∼20–30 μm). Each tank was equipped with a recirculating pump (Hydor Pico 300) and aquarium LED lights (Aqua Illumination Prime 16 HD LED). Lights followed a 12:12 light:dark cycle, featuring two-hour sunrise and sunset ramps (06:00–08:00 and 16:00–18:00) with a peak intensity of ∼100 μmol photons m^−2^ s^−1^.

Within each experimental tank, temperature (°C) and light (lux) was measured every 5 minutes via a HOBO Temperature/Light pendant logger (MX2202; Onset Computer Corp, temperature accuracy ±0.5°C from-20° to 70°C, resolution = 0.04°C, response time = 7 minutes typical to 90% in stirred water; Light accuracy = ±10% typical for direct sunlight). The loggers were cross-calibrated for temperature by placing them all in the same waterbath for 24 hours and for light using a LICOR LI-1500 (LICOR) for 24 hours under 0–200 μmol photons m^−2^ s^−1^ following (Long et al., 2012). Discrete measurements of temperature, pH, salinity and light were taken daily; a digital thermometer (4000EA, Traceable® Products) was used to measure temperature (°C), pH was measured in mV using an InLab Expert Pro pH probe (Cat # 51343101; Mettler Toledo) and salinity was measured in psu on a DuraProbe 4-Electrode Conductivity Cell (Cat # 013005MD, Thermo Fisher Scientific) with pH and salinity probes attached to a Orion Star A series logger (Cat # A325; Thermo Fisher Scientific), and light was measured in μmol photons m^−2^ s^−1^ with an Underwater Quantum Meter (Model MQ-510: Apogee). To quantify pH on the total scale, a calibration curve of mV versus temperature was generated biweekly using Tris standard solution from the Dickson Lab (Batch 127), and total pH was calculated from values of pH (mV) and temperature measured in each tank using the package seacarb (v3.2; Gattuso et al., 2024) in R (v4.4.2) and RStudio (v2024.09.1; R Core Team, 2024). Tanks and corals were cleaned weekly to remove microalgae. Acclimation temperatures and discrete measurements can be found in Figs. S1, S2, and S3.

**Fig. 1.**
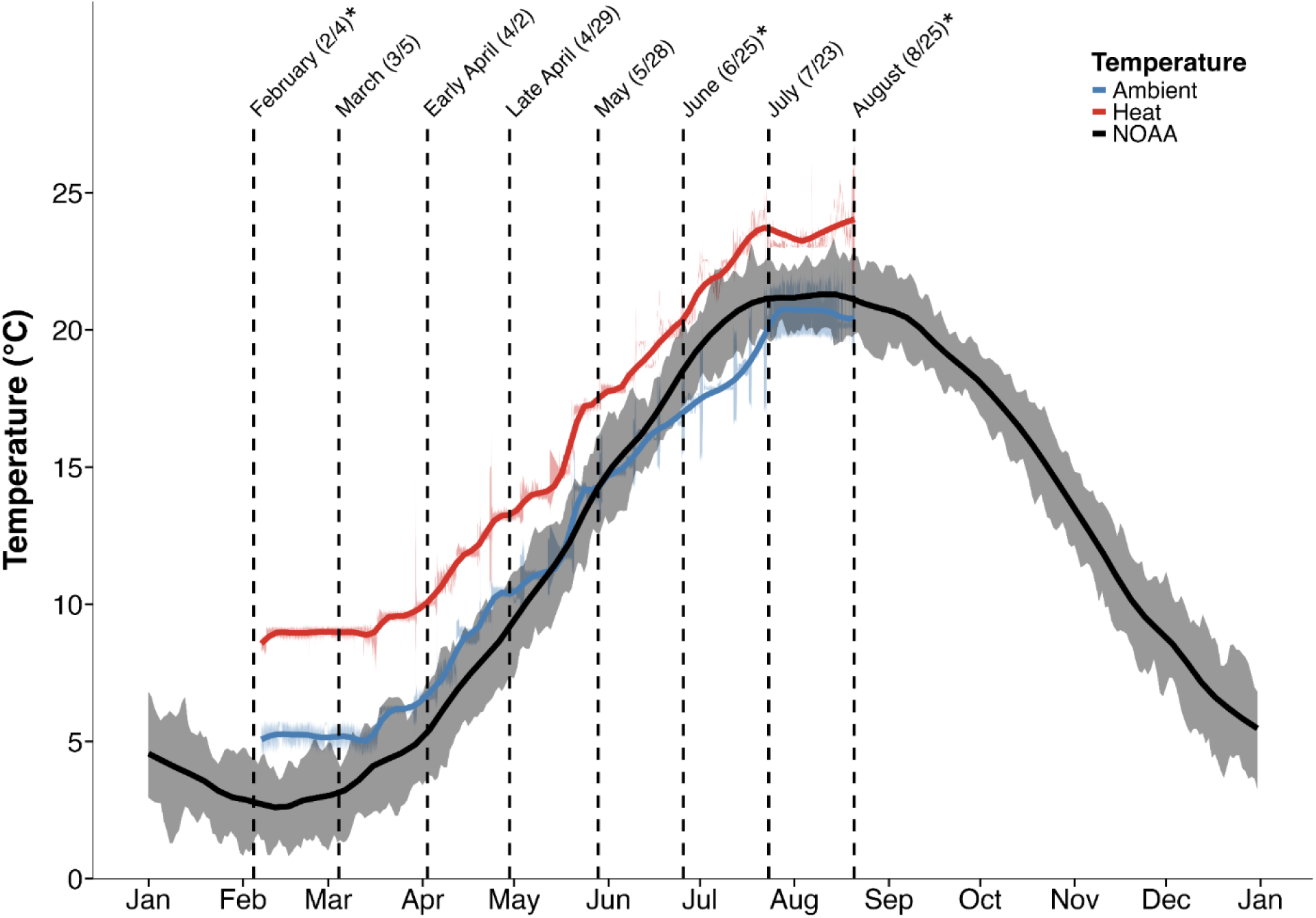
Temperature exposures from February to August 2021. The blue line represents ambient temperature treatment, the red line represents heat temperature treatment, and the black line represents historical average data from the NOAA Station NWPR1 (8452660; Newport, RI). Shading surrounding the experimental treatment lines represents the standard deviation. Vertical dotted lines indicate physiological sampling time points, and asterisks (*) denote time points where samples were collected for molecular analysis (mRNA and miRNA).

**Fig. 2.**
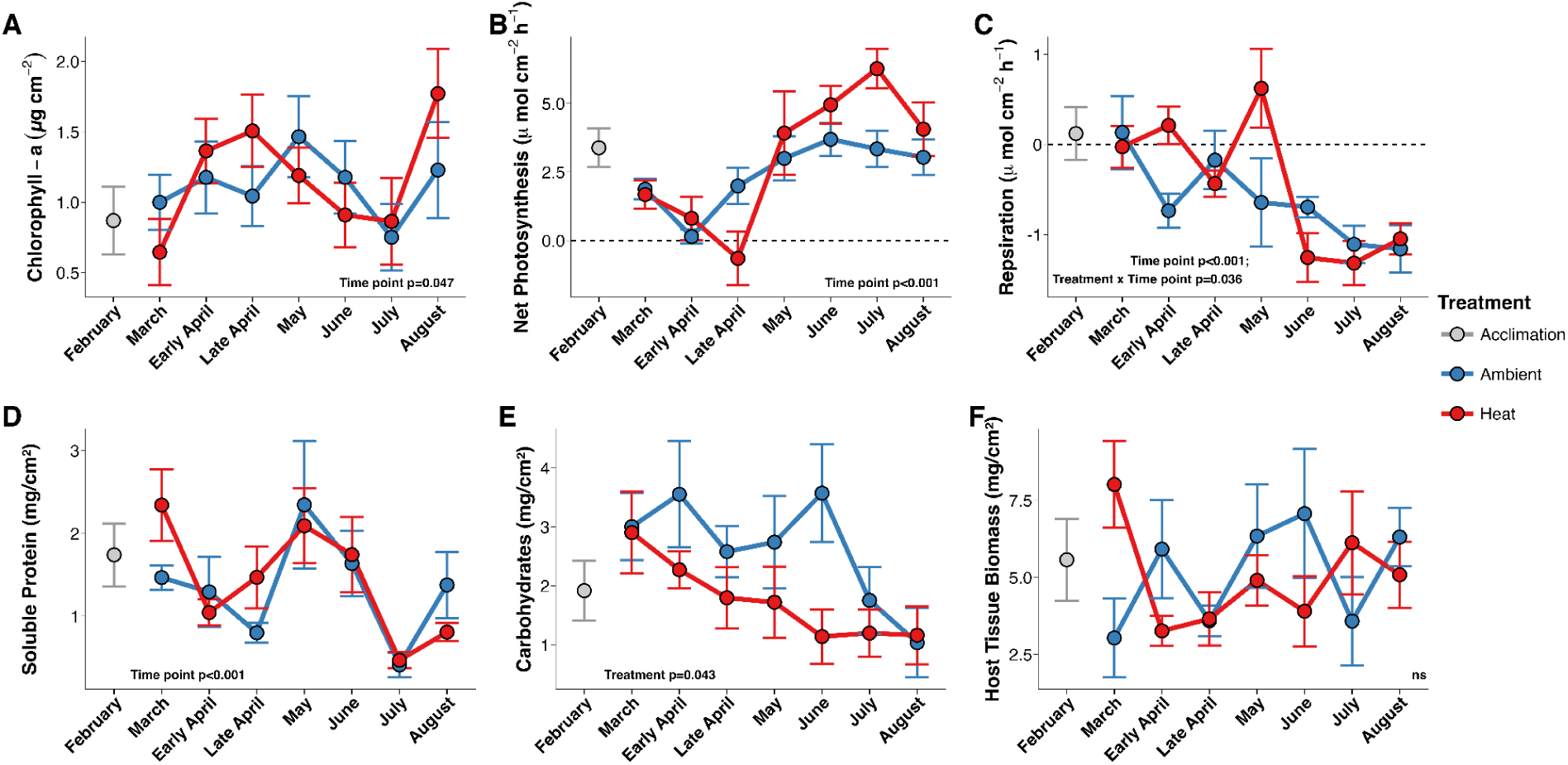
Physiological measurements across the experimental period. A) Chlorophyll-a (µg/cm^-2^), B) Net photosynthesis (µmol O_2_ cm^-2^ h^-1^), C) Respiration (µmol O_2_ cm^-2^ h^-1^), D) Soluble protein (mg cm^-2^), E) Carbohydrates (mg cm^-2^), and F) Host tissue biomass (mg cm^-2^). Data points are colored by treatment (Acclimation: grey; Ambient: blue; Heat: red). Circles represent the mean value at each sampling time point and treatment, with error bars indicating the standard error of the mean (SEM). Full statistical results can be found in Table S1.

Temperature treatments were set based on temperature data from 2005 to 2020 in Newport, RI (Station NWPR1, Newport, RI, NOAA National Ocean Service Water Level Observation Network; 41.504,-71.326). Experimental discrete measurements can be found in Figs. S3 and S4. Corals were randomly allocated to temperature treatments on 1 February 2021, with ambient temperatures starting at 5°C (n=8 tanks) and high temperatures starting at 8°C (n=8 tanks; ambient + 3°C). Experimental temperatures were controlled using a Neptune Apex Aquarium Controller system in combination with a submersible heater (Aqueon Pro Heater, AP300W-H) for high temperatures and an external chiller (AquaEuroUSA Model MC-1/13HP) for ambient temperatures. Every week, temperatures in both treatments were increased by 0.5°C to mimic *in situ* temperature changes (Fig. 1). Samples were collected for metabolic (n=1 per tank, 8 per treatment) and physiological analysis (n=1 per tank, 8 per treatment) on 4 February, 5 March, 2 April, 29 April, 28 May, 25 June, 23 July, and 25 August 2021. Samples were collected for molecular analysis (n=4 per treatment) on 4 February, 25 June, and 25 August 2021 (Fig. 1).

### Physiology

#### Respiration and Photosynthesis

Every month, metabolic rates from corals in each treatment (n=8 per treatment) were measured. Photosynthetic rates (i.e., oxygen production) and respiration rates (i.e., oxygen consumption) were measured using a fiber optic dipping spot O_2_ sensor (Oxygen Dipping Probes DPPSt7, PreSens Precision Sensing GmbH) connected to a Fiber Optic Oxygen Transmitter (OXY-10; PreSens Precision Sensing GmbH) in ∼620 mL cylindrical chambers. Corals were placed in the chambers with a magnetic stir bar and 1µm filtered seawater (FSW) that was pre-heated to the experimental temperature. The chambers were placed on a submersible magnetic stir plate in a water bath heated to the experimental temperatures. Two chambers per stir plate were filled with 1µm FSW to serve as blanks. Photosynthesis rates were measured under a PAR irradiance of ∼400 μmol photons m^−2^ s^−1^ (Aichelman et al., 2019) with light generated from aquarium LED lights (Aqua Illumination Prime 16 HD LED) for 20 minutes. Respiration rates (light enhanced dark respiration, LEDR; Edmunds & Davies, 1988) were measured directly after photosynthetic measurements for 20 minutes in the dark. Following measurements, corals were placed back in their experimental tanks for 24 hours and then sampled. Samples were snap-frozen in liquid nitrogen for subsequent physiological and molecular analyses and stored at-80°C. Rates of oxygen production or consumption were extracted using local linear regressions (alpha=0.2, percentile rank method) in the LoLinR package (v0.0.0.9; Olito et al., 2017) in µmol O_2_ l^-1^ s^-1^.

Rates were corrected to chamber volume including coral displacement and rates from blank chambers were subtracted from the rates of the corals in each run to account for background photosynthesis or respiration owing to phytoplankton, zooplankton, and bacteria. Rates of oxygen production or consumption were normalized to surface area for final units of µmol O2 cm^-2^ h^-1^.

#### Airbrushing and surface area

Prior to airbrushing, small biopsies (∼1 cm^2^) of the snap-frozen coral fragments with tissue on them were clipped from the colony using sterilized clippers and placed in a 1.5 mL tube with 1 mL of DNA/RNA Shield (Cat # R1100, Zymo Research Corp) for molecular analysis. Tissue was removed from the remainder of the snap-frozen colonies using an airbrush (Eclipse Hp-BCS Airbrush 4200, Iwata) with ice cold 1x phosphate buffered saline solution (PBS). The slurry was homogenized using an immersion tip homogenizer (BioGen PRO200 Homogenizer, 10mm probe, PRO Scientific) for one minute at high speed. Following homogenization, aliquots of 1 mL were taken for quantification of soluble protein, carbohydrates, and chlorophyll, while 5 mL was taken for Ash Free Dry Weight (AFDW; tissue biomass). For soluble protein and carbohydrate analysis, host tissue was separated from symbionts and intra-skeletal organisms by centrifuging aliquots at 13,000 g for 3 minutes and using the supernatant for analyses. Surface area following clipping was measured using wax dipping as described in (Veal et al., 2010).

#### Chlorophyll

To isolate symbiont cells for chlorophyll analysis, one mL of tissue homogenate was centrifuged at 13,000 g for 3 minutes and the supernatant was discarded. One mL of 100% acetone was added to the symbiont pellet, vortexed for 30 seconds, and incubated in the dark at 4°C for 24 hours. Samples and acetone blanks were loaded in duplicate (200 µL in each of two wells) into a quartz 96-well plate (Cat# 730-009-44 Hellma Analytics) and read on a plate reader (Synergy HTX Multi-Mode Reader Model S1LFA; BioTek/Agilent) at wavelengths 630, 663, and 750 nm. Chlorophyll-*a* concentrations were corrected for path length (0.584 cm) and calculated using the following equation for dinoflagellates in 100% acetone: Chlorophyll-*a* = (11.43x (E_663 nm_ - E_750 nm_) - 0.64 x (E_630 nm_ - E_750 nm_)) (Jeffrey & Humphrey, 1975). Concentration values were multiplied by 0.584 to correct for the path length of the plate for units µg mL^-1^ and normalized to the airbrushed homogenate volume (mL) and surface area via wax dipping (cm^-2^), for final units of µg cm^-2^.

#### Host Soluble Protein

Host soluble protein was quantified with the Pierce Bicinchoninic Acid (BCA) Protein Assay Kit (Cat # 23225, Pierce Biotechnology) with bovine serum albumin as the standard. Following the manufacturer’s instructions for microplate assay with the generation of a 50:1 of parts A:B working reagent, 25 µL of each sample and standard was added to a 96-well plate in duplicate.

200 µL of the BCA working reagent was added to each well and gently mixed with the sample. The plate was then incubated at 37°C for 30 minutes, cooled to room temperature following incubation, and read on a plate reader (Synergy HTX Multi-Mode Reader Model S1LFA; BioTek/Agilent) at a wavelength of 562 nm. Protein concentrations in µg mL^-1^ were standardized to the airbrushed homogenate volume (mL) and surface area via wax dipping (cm^-2^), for final units of mg cm^-2^.

#### Host Carbohydrates

Host carbohydrate content was quantified using a phenol-sulfuric acid method (Masuko et al., 2005) in a 96-well plate following (Bove & Baumann, 2021). 100 µL of host supernatant was diluted with 900 µL of deionized water. 44 µL of phenol (Cat #A931I-1, Fisher Chemical) and 2.5 mL of sulfuric acid (Cat #A3COSI-200, Fisher Chemical) was added to each sample and also to a series of L-(-)-Glucose standards (0–0.9 mg/mL; Cat #G5500, Sigma Aldrich) and, samples and standards were incubated at room temperature for 30 minutes. Following incubation, 200 µL of each sample and standard in duplicate was loaded into a 96-well plate and read on a plate reader (Synergy HTX Multi-Mode Reader Model S1LFA; BioTek/Agilent) at 485 nm.

Carbohydrate content was normalized to airbrushed homogenate volume (mL) and surface area (cm^-2^), for final units of mg cm^-2^.

#### Host tissue biomass

To quantify AFDW, 5 mL of homogenized tissue slurry was centrifuged at 13000 g for 3 minutes. 4 mL of the host supernatant was removed and placed in a pre-burned aluminum pan weighed on an analytical balance (Mettler Toledo, ML203T/00; Mettler Toledo). Pans were placed in a drying oven (Heraterm General Protocol Oven; Thermo Fisher Scientific) for 24 hours at 60°C, weighed, and then fully dried/combusted in a muffle furnace (Lindberg Blue M Muffle Furnace; Thermo Fisher Scientific) for 4 hours at 450°C. Following this, pans were weighed for a final time and AFDW was calculated as the dry weight minus the ash weight.

AFDW was normalized to airbrushed homogenate volume (mL) and surface area (cm^-2^), for final units of mg cm^-2^.

#### Statistical analysis for physiological variables

For all physiological variables, outliers were identified and removed using the IQR method (Rousseeuw & Croux, 1993). The statistical assumptions of homoscedasticity and of normality of residuals was assessed in the physiological variables using quantile-quantile plots and the Shapiro-Wilk test (Shapiro & Wilk, 1965), respectively. If normality assumptions were violated, the square-root (carbohydrates, AFDW, chlorophyll-a) or inverse (protein) transformations were applied to achieve normality. Two-way ANOVAs were run on all physiological variables with “Treatment” and “Time point” and their interaction as the independent variables (Variable ∼ Treatment*Time point) and post-hoc tests were performed using the Tukey’s Honest Significant Difference (HSD) test (Tukey, 1949), with the TukeyHSD function from the stats package in base R (v4.4.2; R Core Team, 2024).

### Molecular

#### Extractions

DNA and RNA were extracted using the Zymo Quick Miniprep-Plus DNA/RNA kit (CAT D4068, Zymo Research) following the manufacturer’s instructions. Nucleotide quantity and quality were assessed using a Qubit fluorometer (Cat # Q33216, Thermo Fisher/Invitrogen) and 1.5% TAE agarose gel electrophoresis (non-denaturing). mRNA and small RNA libraries were prepared for 20 samples (n=4 per treatment conditions for 5 conditions: February acclimation, June ambient, June heat, August ambient, August heat). mRNA sequencing libraries were prepared using NEBNext Ultra II Directional RNA Library Prep Kit for Illumina following the manufacturer’s instructions (Cat # E7760; New England Biolabs), while small RNA libraries were prepared using NEB Small RNA Library Prep Kit (Cat # E7560; New England Biolabs) following the manufacturer’s instructions. Both the mRNA and small RNA libraries were sequenced on an Illumina HiSeq 4000 in a 2×150bp Paired End (PE) configuration at Genewiz (Azenta Life Sciences). mRNA reads and small RNA reads are stored under NCBI BioProjects PRJNA1231118 and PRJNA1231129, respectively.

#### mRNAs

mRNA sequence quality was assessed before and after trimming using FastQC (v0.11.8; Andrews, 2010) and MultiQC (v1.9; Ewels et al., 2016). Trimming was done using fastp (v0.19.7; Chen et al., 2018) with the following parameters: –detect_adapter_for_pe,–qualified_quality_phred 30, –unqualified_percent_limit 10, –length_required 100, –cut_right cut_right_window 5 cut_right_mean_quality 20. Trimmed reads were aligned to the *A. poculata* reference genome (Stankiewicz et al., 2025) using bowtie2 (v2.4.4; Langmead & Salzberg, 2012). Assembly of reads was performed with stringtie (v2.2.1; Pertea et al., 2015). The stringtie prepDE.py script (Pertea et al., 2015) was then used to create a gene counts matrix from the aligned and assembled data. The counts matrix was filtered to remove any genes that were not expressed across any samples (i.e., 0 gene counts). The pOverA function from the genefilter R package (v1.82.1; Gentleman et al., 2025) was also used to filter for a minimum gene count of 10 in at least 20% of the samples. The resulting gene counts matrix was used as input for expression analysis.

#### miRNAs

miRNA sequence quality was assessed before and after trimming using FastQC (v0.11.8; Andrews, 2010) and MultiQC (v1.9; Ewels et al., 2016). Trimming was done with flexbar (v3.5; Roehr et al., 2017), in which sequences were trimmed to 30 bp, using the following parameters:-a NEB_adapters.fasta,-ap ON,-qf i1.8,-qt 30, –post_trim_length 30. To identify previously described miRNAs, a database containing mature miRNA sequences from miRBase (v22.1; Kozomara et al., 2019) and cnidarian miRNA sequences manually collected from published articles (Ashey et al., 2025) was used as a reference. Previously described and candidate novel miRNAs were identified from alignments of trimmed reads to the *A. poculata* reference genome using ShortStack (v4.0.2; Axtell, 2013; Shahid & Axtell, 2014) with the parameter –dn_mirna, which identifies putative novel miRNAs by searching read-genome alignments for “islands” of significant coverage and evaluating neighboring regions based on a stringent set of miRNA precursor gene characteristics.

#### Differential expression analyses

All gene expression and functional enrichment analyses were performed in R (v4.4.2) and RStudio (v2024.09.1; R Core Team, 2024). Prior to analysis, time point and treatment were combined into a single “Condition” factor (February acclimation, June ambient, June heat, August ambient, and August heat), as recommended for complex experimental designs in the DESeq2 manual (Love et al., 2014). Global expression patterns of mRNAs and miRNAs were first evaluated by Condition using permutational multivariate analysis of variance (PERMANOVA) tests with the adonis2 function in the vegan R package (v2.7-2; Okasen et al., 2017). Significant global differences were further investigated using post-hoc pairwise PERMANOVA tests with the pairwiseAdonis package (v0.4.1; Martinez Arbizu, 2020). For the identification of differentially expressed mRNAs and miRNAs, DESeq2 (v1.40.2; Love et al., 2014) was run with a design of ∼Condition. mRNAs and miRNAs were considered differentially expressed if they had an adjusted p-value < 0.05 and a LFC greater than 1 or less than-1.

#### Seasonal functional enrichment

To characterize the function by season, gene ontology (GO) functional enrichment was first performed on the genes expressed in the February ambient timepoint (relative to the June and August ambient treatments), the June ambient treatment (relative to the February ambient and August ambient treatments), and the August ambient treatment (relative to the February ambient and June ambient treatments). GO enrichment was evaluated with TopGO (v2.58.0; Alexa & Rahnenführer, 2024) using Fisher’s exact test and the weight01 algorithm with a threshold of p<0.05 to assess GO enrichment in Biological Processes in February, June, and August. Unique and shared enriched GO terms were visualized across February, June, and August.

#### mRNA-miRNA interaction prediction and correlation analysis

In cnidarians, miRNAs generally bind with high complementarity to the 3’ untranslated region (3’UTR) or coding sequence (CDS) region of mRNAs, leading to translational repression or degradation (Admoni et al., 2025; Moran et al., 2014). Given the lack of 3’UTR in the *A. poculata* genome, 3’UTRs were operationally defined as the 1000 bp region immediately downstream of the mRNA (Ashey et al., 2025).

Putative mRNA-miRNA interactions were predicted using miRanda (v3.3; Enright et al., 2003) with parameters for strict seed matching (positions 2-8 of the mature miRNA), minimum free energy of duplex formation <-20 kcal/mol, and a minimum alignment score of 120. Both 3’UTRs and CDS sequences were screened as target sites (Moran et al., 2014). Briefly, miRanda determines putative targets using the following criteria: sequence complementarity of mature miRNA to potential target mRNA and the thermodynamic stability of the mRNA:miRNA duplex structure (Enright et al., 2003). To capture the high complementarity binding characteristic of cnidarian miRNA regulation, interactions were required to span at least 13 bp (Admoni et al., 2025; Despard et al., 2025).

To assess condition-specific regulatory relationships, Spearman rank correlations were calculated between miRNA and mRNA expression profiles using vst-normalized counts (Love et al., 2014) within biological replicates of each condition (February acclimation, June ambient, June heat, August ambient, and August heat). To focus on interactions consistent with canonical miRNA-mediated repression, mRNA-miRNA pairs were only retained where the mean miRNA expression exceeded mean mRNA expression, as miRNA regulatory capacity is typically proportional to miRNA abundance relative to target availability (Barbagallo et al., 2023; Denzler et al., 2016; Diener et al., 2024).

#### Network construction and metrics

Unweighted bipartite adjacency networks were constructed for each condition using the predicted interactions identified above. The bipartite structure naturally represents regulatory architecture, with miRNAs and mRNAs forming distinct node classes connected by predicted targeting relationships. For each network, three complementary metrics were calculated to characterize miRNA regulatory properties at each condition: (1) degree–the number of mRNA targets per miRNA, reflecting regulatory breadth; (2) betweenness centrality–the frequency with which a miRNA lies on the shortest path between other miRNAs in the projected network, indicating its role in connecting other regulatory modules; and (3) d’ (specialization index) – a measure of target selectivity, with higher values indicating more specialized miRNA-target relationships (Blüthgen et al., 2006). Networks were compared across seasons (February, June ambient, August ambient) and thermal treatment (June ambient vs. heat, August ambient vs. heat) to assess condition-dependent reorganization of miRNA regulatory networks. The Kruskal-Wallis test (Kruskal & Wallis, 1952) was used to evaluate differences in network metrics across seasons, and the Wilcoxon rank-sum test (Wilcoxon, 1945) was used to evaluate differences in network metrics between treatments in June and August.

#### Correlation trajectory clustering and functional enrichment analysis

To identify mRNA-miRNA regulatory relationships with distinct temporal dynamics across seasons, mRNA-miRNA pairs that were predicted and expressed at all three seasonal timepoints (February, June ambient, August ambient) were retained for analysis. A quadratic polynomial regression model was fitted to the correlation values for each mRNA-miRNA pair across the 3 timepoints, and the regression coefficients were extracted as features capturing the shape of each temporal pattern. K-means clustering (k=5) was applied to group pairs by their correlation direction, enabling identification of mRNA-miRNA pairs with similar regulatory trajectories through time. To connect these temporal regulatory patterns to function, GO enrichment analysis for Biological Processes (BP) terms was performed with TopGO (v2.58.0; Alexa & Rahnenführer, 2024) as detailed above on mRNA targets within each cluster that exhibited negative correlations, consistent with canonical miRNA-mediated repression.

June ambient and heat and August ambient and heat treatments were also evaluated to assess mRNA-miRNA regulatory relationships. For each time point, mRNA-miRNA pairs that were predicted and expressed in both the ambient and heat treatment were retained for analysis. For each pair, a linear regression model was fit, with the regression coefficients extracted as features capturing the shape of the ambient—heat treatment difference. K-means clustering (k=3) was applied to group pairs by their correlation dynamics. GO enrichment analysis was performed with TopGO (v2.58.0; Alexa & Rahnenführer, 2024)) as detailed above on mRNA targets within each cluster that exhibited negative correlations, consistent with canonical miRNA-mediated repression.

## Results

### Physiology

#### Chlorophyll, Photosynthesis, and Respiration

The physiological variables responded much more strongly to the seasonal time points than to the increased temperature treatment at any time point (Fig. 2, Table S1). Chlorophyll-a concentration was significantly different on average by time point (Timepoint, p=0.047), with a pattern of increase through time, but Tukey’s post-hoc test did not identify significant differences between individual time points (Fig. 2A, Table S1).

Net photosynthetic rates increased significantly through time (Timepoint, p<0.001; Fig. 2B, Table S1), but did not differ by treatment (p=0.340), or the interaction of treatment and timepoint (p=0.066). Following initial levels in February, photosynthetic rates declined by approximately 50% to reach a seasonal low in April (Table S1). Rates on average across treatments then increased nearly four-fold from April lows to reach a peak in July (Table S1).

Respiration rates of *A. poculata* were significantly impacted by time point (p<0.001) and the interaction between treatment and time point (p=0.036). Overall, there was not a consistent effect of Treatment (p=0.579; Fig. 2C, Table S1). Respiration remained low throughout the spring before increasing significantly to reach peak levels on average across treatment in July and August (Table S1). While there were no significant differences in respiration between treatments within any single time point (p > 0.05), the interaction was driven by divergent seasonal trajectories of the two treatment groups. Heat-treated colonies experienced the lowest respiration rates in May before undergoing a sharp four-fold increase to reach a peak in June and July (Table S1). In contrast, colonies in the ambient treatment had a more gradual increase in respiration rates over the same period (Fig. 2C). By July and August, respiration in both treatments converged at their highest observed levels, representing a significant increase in metabolic demand compared to the initial spring months (Table S1).

#### Host Soluble Protein, Carbohydrates and Biomass

Soluble protein content of *A. poculata* was not affected by treatment (p=0.995) or the interaction between time point and treatment (p=0.574), but exhibited significant seasonal shifts (p<0.001; Fig. 2D, Table S1). For the majority of the study period (February–June), protein levels were relatively stable (Fig. 2D). However, protein content decreased by approximately 75% in July, reaching a seasonal minimum that was significantly lower than all other time points (p < 0.05; Table S1). Following this mid-summer low, protein levels began to recover in August, increasing nearly three-fold (Fig. 2D; Table S1).

Carbohydrate content was marginally significant by treatment (p=0.043), and the post-hoc test revealed that the ambient treatment had higher values than the high treatment (p=0.043; Fig. 2E, Table S1); however, carbohydrate content was not significantly different by time point (p=0.094) or the interaction term (p=0.747). Tissue biomass was not significantly different by treatment (p=0.825), time point (p=0.729), or the interaction term (p=0.102; Fig. 2F, Table S1).

### Molecular

#### mRNAs

mRNA-sequencing yielded an average of 33,916,757 raw reads per sample (Table S2). Samples retained an average of 22,592,186 reads after trimming (Table S2). Alignment with bowtie2 to the *A. poculata* reference genome resulted in an average alignment rate of 36.2% (Table S2), which is comparable to previous studies with this species (Borbee et al., 2026; Wuitchik et al., 2021, 2024).

#### miRNAs

Small RNA-sequencing yielded an average of 19,491,009 raw reads per sample (Table S2). After trimming, samples retained an average of 18,033,790 reads (Table S2). 51 putative miRNAs were identified using ShortStack: five known and 46 novel (Table S3). The known miRNAs corresponded to those identified in previous cnidarian studies: miR-2030, miR-2037, miR-9425, miR-2036, and miR-100.

#### Differential expression of DEGs and DEMs

After filtering for lowly expressed genes, 28,720 genes remained for downstream expression analysis out of 47,156 total genes. Global mRNA expression patterns exhibited significant differences by condition (PERMANOVA p=0.033; Table S4), driven by the transition from the February baseline to later time points (p<0.04 for all February contrasts; Table S5). When examining shifts only for the experimental period (i.e., June and August), no significant differences were detected between the summer contrasts (PERMANOVA p=0.358; p>0.1 for all June and August contrasts; Tables S4 and S5).

Differential expression analysis between time points under ambient conditions revealed 666 DEGs between February and June ambient, with 514 upregulated in February and 152 upregulated in June ambient (Table S6). Between February and August ambient, 671 genes were differentially expressed, 449 upregulated in February and 222 upregulated in August ambient (Table S6). Comparisons between ambient June and August timepoints showed 176 DEGs, with 53 upregulated in June ambient and 123 upregulated in August ambient (Table S6). In line with the lack of strong separation in global expression between the temperature treatments in June and August, there were also fewer DEGs identified between treatment comparisons (Table S6). June ambient and heat treatments had 98 DEGs, with 44 upregulated under June ambient conditions and 54 upregulated under June heat conditions (Table S6). While the number of DEGs remained low in August (62 DEGs), only 15 were upregulated in August ambient conditions, whereas 47 were upregulated in August heat conditions (Table S6). PCA of the DEGs found that the February samples clustered away from the June and August ambient and heat treatments along the PC1 axis (Fig. 3A), accounting for 33% variance.

**Fig. 3.**
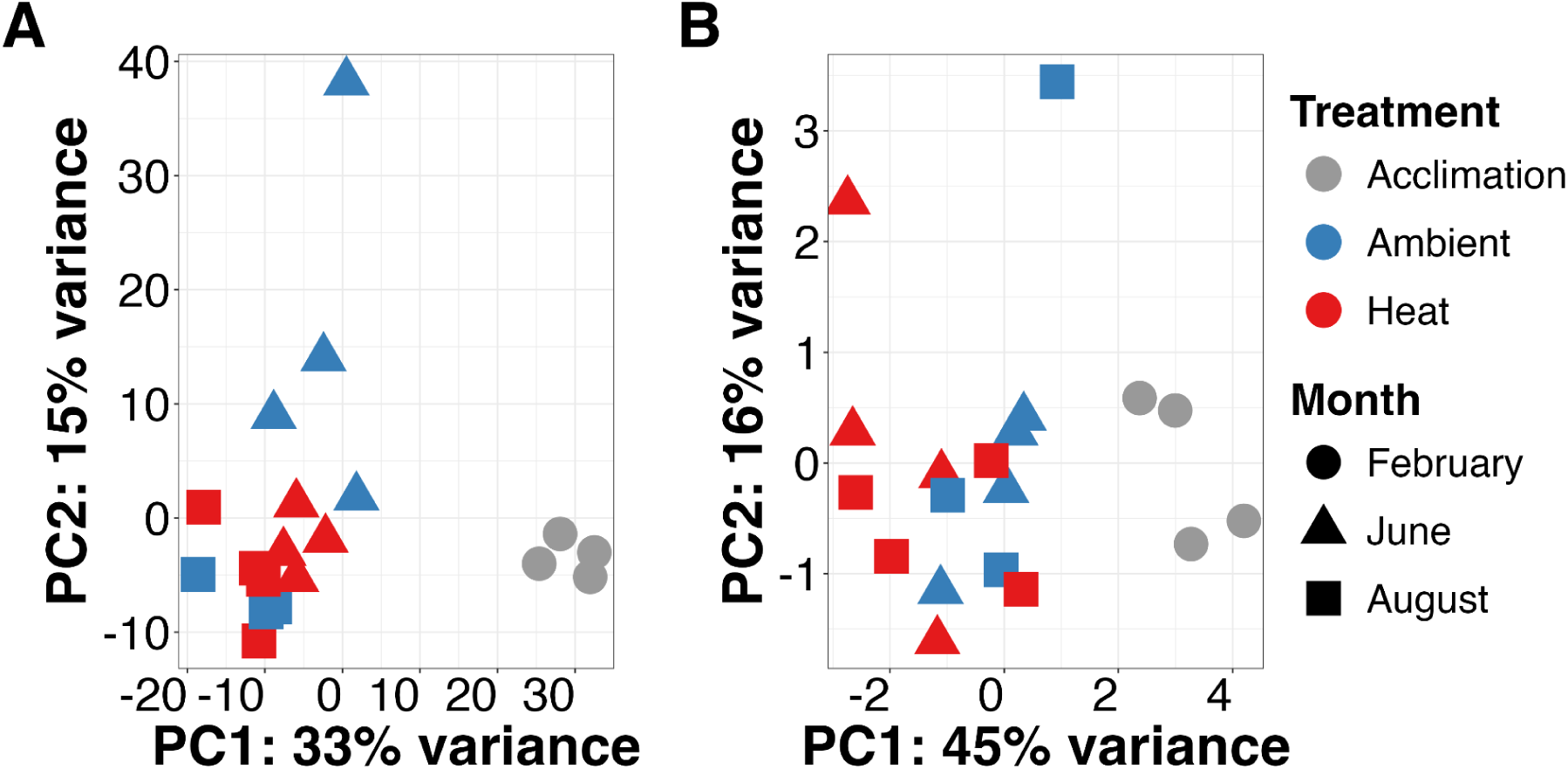
Principal component analyses of (A) differentially expressed mRNAs and (B) differentially expressed miRNAs. Each point represents an individual coral colony. Samples are color-coded by treatment (Acclimation: grey; Ambient: blue; Heat: red) and shared by time point (February: circles; June: triangles; August: squares). The first principal component (PC1) represents the largest proportion of total variance for both mRNA (33%) and miRNA (45%) datasets.

Similarly to the mRNA results, global miRNA expression patterns differed significantly by condition (PERMANOVA p=0.006; Table S4), a trend primarily driven by the separation of February and all later time points (p<0.04 for all February contrasts, except August heat; Table S5). While a global shift in miRNA expression persisted even after the removal of February (PERMANOVA p=0.018; Table S4), post-hoc comparisons did not find any significant differences between the summer conditions (p>0.08 for all June and August contrasts; Table S5). Comparison of DEMs between time points under ambient conditions revealed two DEMs between February and June ambient, one upregulated in February and one upregulated in June (Table S7). Between February and August ambient, two DEMs, miR-100 and Cluster_1453, were upregulated in February. No DEMs were identified between June and August ambient.

There were no DEMs between the June ambient and heat treatments or the August ambient and heat treatments. Similarly to the PCA of DEGs, the PCA of the DEMs (Fig. 3B) demonstrated that February samples clustered away from the June and August ambient and heat treatments along the PC1 axis, accounting for 45% variance.

#### Seasonal functional enrichment of expressed mRNAs

Genes expressed in February, relative to the June and August ambient treatments, were significantly enriched in 114 BP GO terms, 103 of which were unique to February (Table S8; Fig. 4A). The top ten unique terms (in terms of p-value significance) were positive regulation of fibroblast growth factor receptor, regulation of complement activation, Mo-molybdopterin cofactor biosynthetic process, complement activation classical pathway, spermatogenesis, protein quality control for misfolded or incomplete proteins, vascular endothelial growth factor receptor signaling, regulation of macrophage chemotaxis, Hsp90 deacetylation, and polyubiquitinated misfolded protein transport (Fig. 4B).

**Fig. 4.**
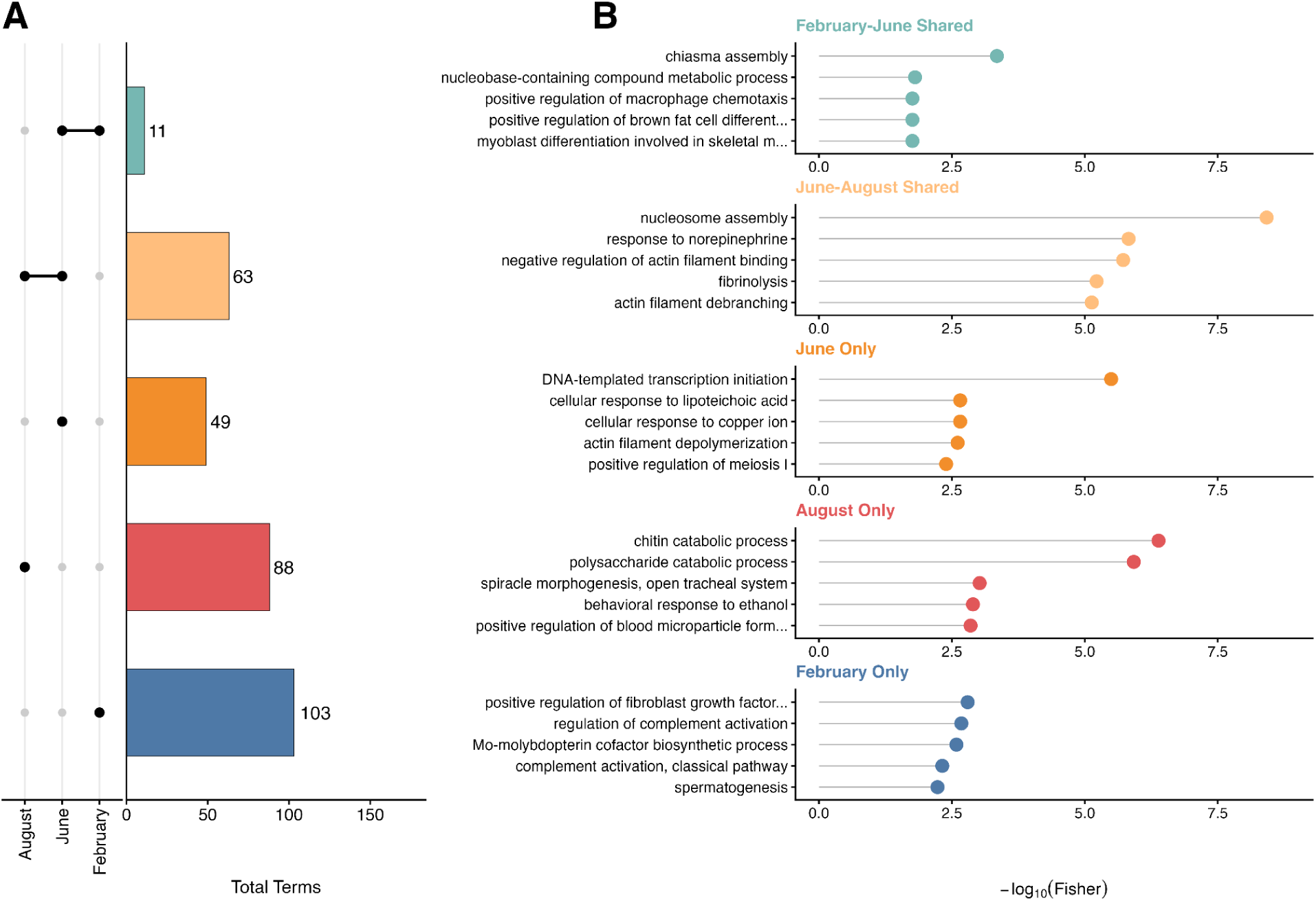
Comparative enrichment analysis of seasonal transcriptomic shifts. A) UpSet plot displaying the distribution of enriched Biological Processes (BP) gene ontology (GO) terms across February, June, and August ambient conditions. Vertical bars indicate the count of enriched terms unique to a specific time point or shared across time points. GO terms that are shared between time points are denoted by connected black dots below the x-axis. B) The top five GO terms (by p-value) for each intersection or unique set are shown to the right of the corresponding bar.

Genes expressed in the June ambient treatment, relative to the February acclimation and August ambient treatments, were significantly enriched in 123 BP GO terms, 49 of which were unique to June (Table S8; Fig. 4A). The top ten unique terms (in terms of p-value significance) were DNA-templated transcription initiation, cellular response to copper ion, cellular response to lipoteichoic acid, actin filament depolymerization, positive regulation of meiosis I, positive regulation of reciprocal meiotic recombination, notochord development, meiotic gene conversion, cellular response to histamine, microtubule-based process (Fig. 4B).

Finally, genes expressed in the August ambient treatment, relative to the February acclimation and June ambient treatments, were significantly enriched in 151 GO terms, 88 of which were unique to August only (Table S8; Fig. 4A). August enriched terms included immune system process, chitin catabolic process, polysaccharide catabolic process, response to ethanol positive regulation of blood microparticle formation, positive regulation of cholesterol storage, positive regulation of macrophage derived foam cell differentiation, regulation of catalytic activity, extracellular matrix organization, and protein peptidyl-prolyl isomerization (Fig. 4B).

No significantly enriched terms were shared across February, June, and August, nor were enriched terms shared only between February and August (Fig. 4A). 11 enriched terms were shared between February and June: myoblast differentiation involved in skeletal muscle, positive regulation of brown fat cell differentiation, positive regulation of macrophage chemotaxis, positive regulation of myoblast differentiation, positive regulation of myoblast fusion, positive regulation of myotube differentiation, stress-induced premature senescence, cell surface receptor protein serine/threonine kinase activity, chiasma assembly, xylulose metabolic process, and nucleobase-containing compound metabolic process (Fig. 4A, B). The summer months, June and August, had 63 unique enriched GO terms shared between them (Fig. 4A). These included skeletal system development, nucleosome assembly, blood vessel development, collagen fibril organization, positive regulation of ERK1 and ERK2 cascade, transforming growth factor beta receptor signaling, fibrinolysis, muscle tissue development, cellular response to amino acid stimulus, and Rho protein signal transduction, among others (Fig. 4B, Table S8).

#### Treatment functional enrichment of expressed mRNAs June and August

There was significant enrichment from the DEGs between the 4 treatment comparisons (June ambient, June high, August ambient, August High). In June, the ambient treatment exhibited enrichment in 31 BP GO terms relative to the heat treatment. GO Processes involving multiple significant genes within a single term included multicellular organism development, microtubule-based processes, and nucleosome assembly (Table S9), and the rest were driven by single genes. From the 27 BP terms in the June heat treatment, GO terms of cobalt ion and cobalamin transport were enriched. Positive regulation of protein homodimerization activity was significantly enriched in both June treatments. DEGs from August ambient treatment relative to the August heat treatment were characterized by 61 enriched BP terms, although only the term defense response to bacterium was associated with more than one gene (Table S9). Under the August heat treatment, 39 BP terms were enriched, with glucose metabolic process and DNA replication being the only terms represented by multiple genes (Table S9). The August treatments shared a single overlapping term: cellular response to lipoteichoic acid.

#### Network analysis of mRNA-miRNA interactions

Analysis of network topology of the mRNA-miRNA interactions (Table S10) across the three seasonal ambient conditions revealed considerable stability. No significant differences were detected in miRNA degree (regulatory breadth; p=0.711), betweenness (connectivity; p=0.910), or specialization (target selectivity; p=0.800) across February, June, and August. Similarly, no significant differences were detected in miRNA degree (June Ambient vs. Heat p=0.997; August Ambient vs. Heat p=0.803), betweenness (June Ambient vs. Heat p=0.946; August Ambient vs. Heat p=0.881), or specialization index (June Ambient vs. Heat p=0.934; August Ambient vs.

Heat p=0.933) between treatments at either June or August time points.

#### Trajectory analysis of seasonal mRNA-miRNA interactions

A total of 36,925 predicted mRNA-miRNA interactions spanning February, June ambient, and August ambient were identified between 51 miRNAs and 8,950 mRNAs. Expression-based filtering to retain only interactions where mean miRNA expression exceeded mean mRNA expression–consistent with miRNA repression of mRNA–reduced the dataset to 22,994 interactions across conditions remained, representing 49 miRNAs and 7,056 mRNAs. The number of interactions per condition were: 6,683 in February; 7,927 in June; and 7,684 in August. Of those, 5,785 mRNA-miRNA interactions were consistent across the three seasonal timepoints (February, June ambient, August ambient). K-means clustering of the quadratic regression coefficients grouped these pairs into five distinct temporal correlation clusters (Fig. 5), representing different regulatory dynamics over time.

**Fig. 5.**
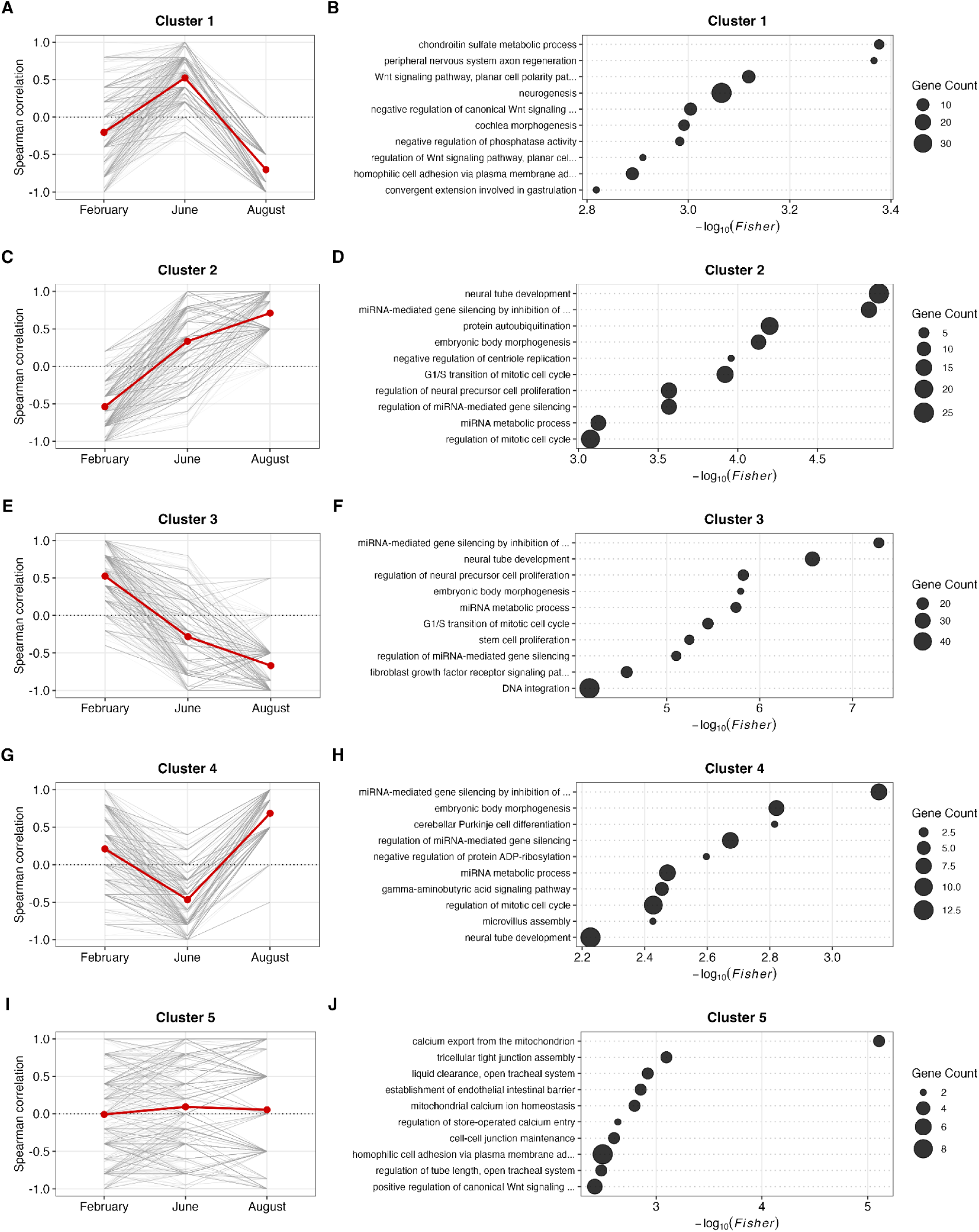
Seasonal correlation trajectories of mRNA-miRNA pairs across three time points (February, June, August). (A, C, E, G, I) Individual pair trajectories are represented by grey lines, while the mean of pair trajectories is highlighted by the red line. (B, D, F, H, J). Dot plots illustrating the significant enrichment of the top ten (by p-value) Biological Processes (BP) GO terms. The x-axis represents the significance value, the y-axis represents the specific terms, and the size of each dot corresponds to the gene count associated with the specified biological process. See Table S11 for all enriched terms.

To assess the biological functions of mRNAs that were putatively targeted and repressed by miRNAs within each cluster, we examined the negative correlations (i.e., high expression miRNA, low expression mRNA). Cluster 1 exhibited an inverted U shape with mean negative mRNA-miRNA correlations in February and August (i.e., repressed) (Fig. 5A). Significantly enriched terms (n=143 BP) in the putatively targeted and repressed mRNAs were related to development and growth including peripheral nervous system axon regeneration, Wnt signaling pathway with planar cell polarity patterning; neurogenesis, negative regulation of canonical Wnt signaling pathway, cochlea morphogenesis, regulation of Wnt signaling pathway (planar cell polarity pathway), convergent extension involved in gastrulation, chondroitin sulfate metabolic process, negative regulation of phosphatase activity, and homophilic cell adhesion via plasma membrane adhesion (Fig. 5B; Table S11).

Cluster 2 showed mean negative correlations occurred in February (Fig. 5C). Enriched GO terms (139 significantly enriched BP GO terms) among negatively correlated targets (i.e., February) included neural tube development, miRNA-mediated gene silencing by inhibition of translation, protein autoubiquitination, embryonic body morphogenesis, negative regulation of centriole replication, G1/S transition of mitotic cell cycle, regulation of neural precursor cell proliferation, regulation of miRNA-mediated gene silencing, miRNA metabolic process, and regulation of mitotic cycle (Fig. 5D; Table S11).

Cluster 3 displayed mean negative correlations emerging in June and more strongly in August (Fig. 5E). There were 82 significantly enriched BP GO terms among negatively correlated mRNA targets. GO enrichment identified significant terms including neural tube development, miRNA-mediated gene silencing by inhibition of translation, regulation of neural precursor cell proliferation, embryonic body morphogenesis, miRNA metabolic process, G1/S transition of mitotic cell cycle, stem cell proliferation, regulation of miRNA-mediated gene silencing, fibroblast growth factor receptor signaling patterns, and DNA integration (Fig. 5F; Table S11).

Cluster 4 exhibited a U shape, with mean negative correlations in June, and 78 significantly enriched BP GO terms (Fig. 5G). Negatively correlated targets in this cluster were enriched for miRNA-mediated gene silencing by inhibition of translation, embryonic body morphogenesis, cerebellar Purkinje cell differentiation, regulation of miRNA-mediated gene silencing, negative regulation of protein ADP-ribosylation, miRNA metabolic process, gamma-aminobutyric acid (GABA) signaling pathway, regulation of mitotic cell cycle, microvillus assembly, and neural tube development (Fig. 5H; Table S11).

Cluster 5 represented a distinct subset of mRNA-miRNA pairs maintaining consistent correlation patterns over time, indicating stable regulatory relationships across seasons (Fig. 5I). 107 BP GO terms were significantly enriched in this cluster. When examining the negatively correlated targets in this cluster (i.e., all lines fully below 0), terms including calcium export from the mitochondrion, tricellular tight junction assembly, liquid clearance in open tracheal system, establishment of endothelial intestinal barrier, mitochondrial calcium ion homeostasis, regulation of store-operated calcium entry, cell-cell junction maintenance, homophilic cell adhesion via plasma membrane adhesion, regulation of tube length in open tracheal system, and positive regulation of canonical Wnt signaling pathway were enriched (Fig. 5J; Table S11).

#### Trajectory analysis of June treatment mRNA-miRNA interactions

K-means clustering of the 7,075 mRNA-miRNA pairs in the June treatments grouped these pairs into three distinct correlation clusters, representing different regulatory dynamics across the treatments.

To assess the biological functions of mRNAs that were putatively targeted and repressed by miRNAs, we examined the negative correlations of each cluster. June Cluster 1 exhibited an increasing pattern, with mean negative mRNA-miRNA correlations in the June ambient treatment and mean positive correlations in the June heat treatment (Fig. 6A). GO enrichment of negatively correlated targets revealed 132 BP GO terms significantly enriched. Cilium-dependent cell motility, cell development, BMP signaling pathway, MDA-5 signaling pathway, protein polyubiquitination, intracellular copper ion homeostasis, axial mesoderm formation, lung alveolus development, resolution of mitotic recombination intermediates, and response to cholesterol were among the top enriched terms (Fig. 6B, Table S12).

**Fig. 6.**
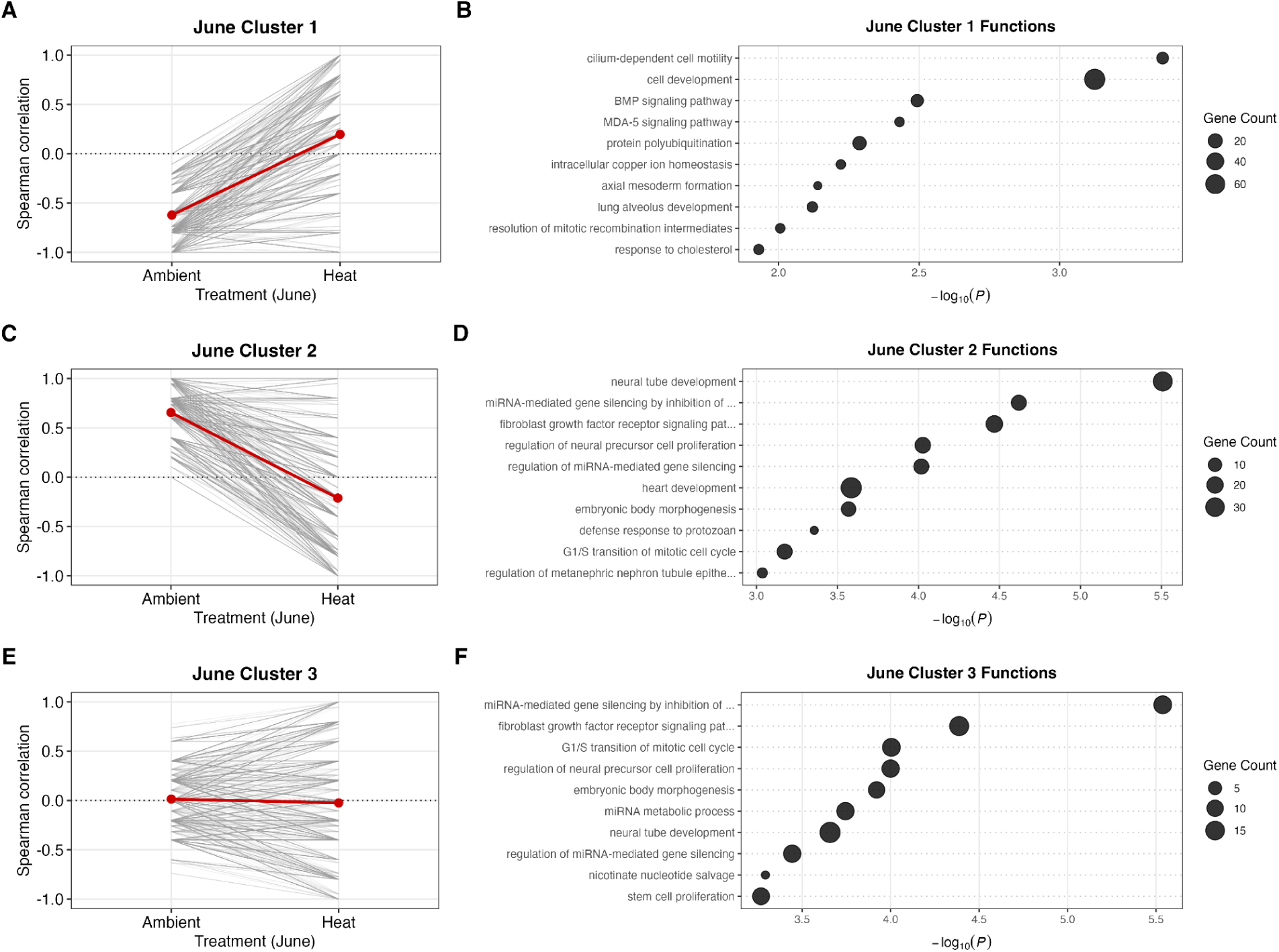
June clusters and enriched terms. June treatment correlation trajectories of mRNA-miRNA pairs between Ambient and High treatments. (A, C, E) Individual pair trajectories are represented by grey lines, while the mean of pair trajectories is highlighted by the red line. (B, D, F). Dot plots illustrating the significant enrichment of the top ten (by p-value) Biological Processes (BP) GO terms. The x-axis represents the significance value, the y-axis represents the specific terms, and the size of each dot corresponds to the gene count associated with the specified biological process. See Table S12 for all enriched terms.

June Cluster 2 had a decreasing pattern, with mean positive correlations in the June ambient treatment and mean negative mRNA-miRNA correlations in the June heat treatment, (Fig. 6B). Enriched GO terms (158 significantly enriched BP GO terms) among negatively correlated targets included neural tube development, miRNA-mediated gene silencing by inhibition of translation, fibroblast growth factor receptor signaling pathway, regulation of neural precursor cell proliferation, regulation of miRNA-mediated gene silencing, heart development, embryonic body morphogenesis, defense response to protozoan, G1/S transition of mitotic cell cycle, and regulation of metanephric nephron tube epithelial cell differentiation (Fig. 6D, Table S12).

June Cluster 3 represented a distinct subset of mRNA-miRNA pairs maintaining relatively consistent correlation patterns over the two treatments in June, indicating stable regulatory relationships (Fig. 6E). 165 BP GO terms from the correlation values below 0 were significantly enriched in this cluster. Terms included miRNA-mediated gene silencing by inhibition of translation, fibroblast growth factor receptor signaling pathway, G1/S transition of mitotic cell cycle, regulation of neural precursor cell proliferation, embryonic body morphogenesis, miRNA metabolic process, neural tube development, regulation of miRNA-mediated gene silencing, nicotinate nucleotide salvage, and stem cell proliferation (Fig. 6F, Table S12).

#### Trajectory analysis of August treatment mRNA-miRNA interactions

K-means clustering of the 7,038 mRNA-miRNA pairs in the August treatments into three distinct correlation patterns/clusters, representing different regulatory dynamics across the treatments.

To assess the biological functions of mRNAs that were putatively targeted and repressed by miRNAs, we examined the negative correlations of each cluster. August Cluster 1 represented a distinct subset of mRNA-miRNA pairs maintaining relatively consistent correlation patterns over the two treatments in June, indicating stable regulatory relationships (Fig. 7A). 179 BP GO terms were significantly enriched in this cluster. Terms included heterophilic cell adhesion, regulation of platelet activation, calcium-dependent cell adhesion, regulation of leukocyte migration, regulation of integrin activation, leukocyte tethering or rolling, pre-miRNA export from nucleus, microtubule cytoskeletal organization, equator specification, and division septum site selection (Fig. 7B; Table S12).

**Fig. 7.**
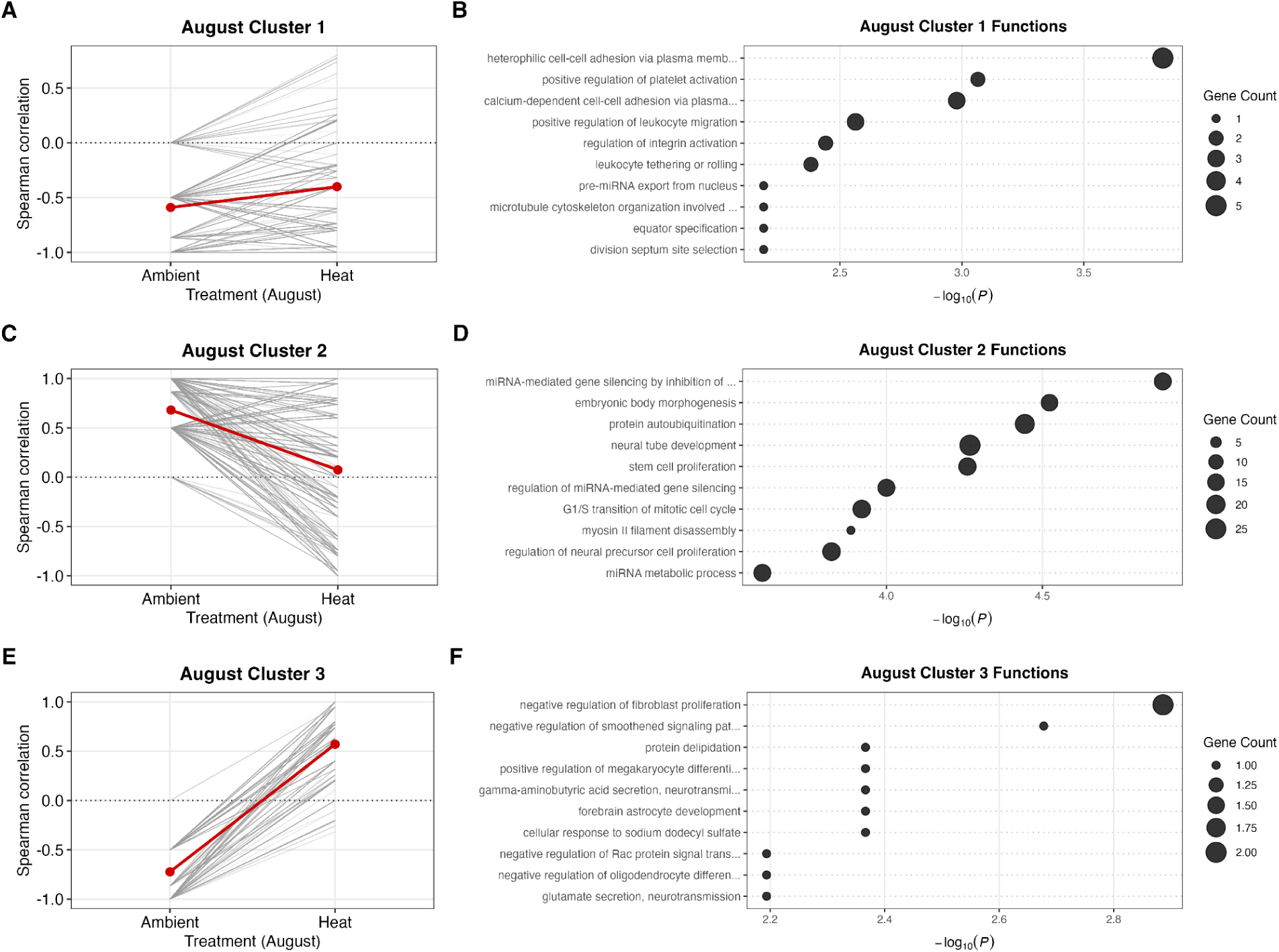
August clusters and enriched terms. Treatment correlation trajectories of mRNA-miRNA pairs between Ambient and High treatments. (A, C, E) Individual pair trajectories are represented by grey lines, while the mean of pair trajectories is highlighted by the red line. (B, D, F). Dot plots illustrating the significant enrichment of the top ten (by p-value) Biological Processes (BP) GO terms. The x-axis represents the significance value, the y-axis represents the specific terms, and the size of each dot corresponds to the gene count associated with the specified biological process. See Table S12 for all enriched terms.

August Cluster 2 demonstrated a downward pattern between ambient and high treatments in August, with mean positive correlations in the ambient treatment and mean negative mRNA-miRNA correlations in the heat treatment, with (Fig. 7C). Enriched GO terms (152 significantly enriched BP GO terms) among negatively correlated targets included miRNA-mediated gene silencing, embryonic body morphogenesis, protein autoubiquitination, neural tube development, stem cell proliferation, regulation of miRNA-mediated gene silencing, G1/S transition of mitotic cell cycle, myosin II filament disassembly, regulation of neural cell proliferation, and miRNA metabolic process (Fig. 7D; Table S12).

August Cluster 3 exhibited an upward pattern between ambient and high treatments in August with mean negative mRNA-miRNA correlations in the ambient treatment, with mean positive correlations in the heat treatment (Fig. 7E). GO enrichment of negatively correlated targets revealed 171 BP GO terms significantly enriched. Terms including regulation of fibroblast proliferation, regulation of smoothed signaling pathways, protein delipidation, regulation of megakaryocyte differentiation, gamma-aminobutyric acid secretion, forebrain astrocyte development, response to sodium dodecyl sulfate, regulation of Rac protein signal transduction, regulation of oligodendrocyte differentiation, and glutamate secretion were enriched (Fig. 7F; Table S12).

## Discussion

While corals are acutely threatened by climate change, the molecular mechanisms governing their acclimatory plasticity remain underdescribed. The temperate coral, *Astrangia poculata*, provides a unique system for investigating these mechanisms; it routinely endures an extreme range of seasonal temperatures (-1°C to 27°C) at the northern edge of its distribution (Ashey et al., 2025; Jacques et al., 1983), a region where seawater temperatures have already increased substantially and are continuing to warm (Ahmed et al., 2025; Young & Young, 2025). Here, we demonstrate that season acts as the primary driver of both physiology and gene expression, and miRNA-mediated regulation contributes to the physiology of winter quiescence and summer metabolic and reproduction activity. Furthermore, the miRNA-mRNA regulatory program appears to be robust to the moderate thermal stress of +3°C across the seasons, enabling controlled and responsive transcriptional activity in this eurythermal coral.

### Seasonality is the primary driver of physiology and gene expression

Consistent with previous studies of Rhode Island *A. poculata*, seasonal environmental conditions are primary drivers of physiological differences (Brown et al., 2022; Dimond & Carrington, 2007, 2008; Sharp et al., 2017; Trumbauer et al., 2021, 2022). In our study, we observed that photosynthesis and respiration were lowest in colder winter months, reflecting a state of metabolic dormancy (Grace, 2017). This is consistent with prior findings (Jacques et al., 1983), which demonstrated that both respiration and photosynthesis are severely depressed at ∼6°C. These rates subsequently rose in tandem with temperature as chlorophyll-a increased, mirroring established patterns where chlorophyll levels increase as ambient temperatures rise (Jacques et al., 1983; Scheufen et al., 2017). Physiological differences between the ambient and the 3°C heated treatments were limited in contrast with these robust seasonal physiological shifts.

Physiologically, the largest change across a timepoint that we documented was in soluble protein content from June to July. Proteins are typically catabolized after carbohydrate and lipid reserves have already been depleted, which can happen under thermal stress bleaching and starvation (Rodrigues & Grottoli, 2007). However, in our study, carbohydrate levels and biomass remained stable, suggesting targeted resource reallocation rather than a general depletion of energy stores.

*A. poculata* develops gametes from March to July and typically spawn July-September (Szmant-Froelich et al., 1980). While elevated protein catabolism in *A. poculata* could be utilized to meet nitrogen demands associated with gametogenesis (Jaffe et al., 2023; Szmant et al., 1990), protein rich components are also key to gametogenesis (Timmins-Schiffman et al., 2024) and would be allocated specifically to oocytes (e.g., maternal proteins, mRNAs), and thus would decline rapidly and substantially following a spawning event. Based on histological tracking (Gilligan et al. in prep), spawning in 2021 occurred in July. Therefore, our results support gamete tissue loss via spawning as a driver of protein loss and not catabolism for energetic demands. A trend for protein loss following spawning is seen in tropical coral taxa (Leuzinger et al., 2003), although declines in protein content are generally less pronounced and less consistent than those observed for lipids, but we did not assess lipids in our study.

Seasonal phenotypic plasticity was also strongly present at the transcriptomic level. Our study provides the first characterization of the *A. poculata* transcriptome across seasons, characterizing how temperate corals navigate extreme environmental variability at the molecular level. In our earliest seasonal timepoint, February, water temperatures were ∼4–6°C; at these temperatures, metabolic quiescence of *A. poculata* is common (Jacques et al., 1983; Grace 2017). Gene expression related to protein quality control, misfolded protein management, and cellular stress anticipation indicate that *A. poculata* is prioritizing cellular integrity during winter, preventing cumulative damage when metabolic inputs are reduced (Marescal & Cheeseman, 2020). Similar molecular strategies have been observed in *A. poculata* and *Acropora muricata* exposed to cold stress (6°C and 15°C, respectively), where cellular maintenance was prioritized through the upregulation of proteasomes, translation initiation factors, and actin-related structure proteins (Lee et al., 2018; Wuitchik et al., 2021). Furthermore, prioritizing genome stability and protein quality control during periods of low metabolic activity is a conserved strategy shared widely across other seasonally quiescent organisms (Leung et al., 2024; Rosswag De Souza et al., 2025; Stankiewicz et al., 2025; Waterworth et al., 2015, 2016).

Across the summer months, as temperatures warmed, functional enrichment shifted from cellular maintenance to growth, reproduction, and immune activity. Early summer (June) was characterized by transcriptional signatures of meiotic activity, cytoskeletal remodeling, and environmental sensing–processes consistent with the timing of gametogenesis in *A. poculata* (Szmant-Froelich et al., 1980) and an increase in temperature or physical environmental change (Dimond & Carrington, 2007; Dimond et al., 2013). Historical histological evidence of gametogenesis indicates oogenesis starts in March-April and spawning occurs some time between July–September (Szmant-Froelich et al., 1980) and gametogenesis in scleractinian corals is cued by environmental factors such as photoperiod and seawater temperature (Tan et al., 2025). By late summer (August), enrichment of functions shifted toward heightened immune and metabolic activity, likely reflecting increased microbial pressures and changing host-symbiont-microbiome dynamics during the warmest part of the year (Dimond & Carrington, 2007, 2008; Sharp et al., 2017; Staroscik & Smith, 2004). June and August shared a suite of enriched terms related to skeletal development, cytoskeletal organization, and immune signaling, indicating that summer was broadly defined by dynamic growth (Dellaert et al., 2022; Jacques et al., 1983; Jacques & Pilson, 1980) and bioenergetic flexibility (Lindsay et al., 2025; Trumbauer et al., 2021), even as immune priorities escalated with seasonal warming and associated shifts in host-microbiome dynamics (Sharp et al., 2017).

#### miRNAs facilitate biological requirements of homeostasis, protection and repair through dynamic seasonal regulatory interactions within a stable infrastructure

We found that the miRNA repertoire of *A. poculata* contains five documented and 46 novel miRNAs. This is comparable to other scleractinian corals, which range from 26 to 65 identified miRNAs across various species (Ashey et al., 2025; Despard et al., 2025; Gajigan & Conaco, 2017; Liew et al., 2014; Praher et al., 2021). The five previously documented miRNAs found here are highly conserved across diverse coral lineages (Ashey et al., 2025; Praher et al., 2021). The dynamics of miRNA expression across time points indicate an environmentally responsive role for this regulatory mechanism (Biggar & Storey, 2015; Leung & Sharp, 2007, 2010), in the broad conditions *A. poculata* inhabits across a year (Ashey et al., 2025).

As the first study to characterize miRNAs as seasonally responsive in corals, these findings represent a major advancement in our understanding of how temperate corals generate phenotypic plasticity at the molecular level to navigate extreme environmental variability. Strikingly, the miRNA regulatory system of *A. poculata* appears to be achieved through a juxtaposition of the miRNA-mRNA network architectural stability, yet interaction flexibility. The miRNA-mRNA network topology metrics–degree, betweenness, and target specialization–remained remarkably stable across seasons, while mRNA-miRNA interactions (i.e., correlation coefficients, Fig. 5) were highly dynamic. This suggests that *A. poculata* maintains robust miRNA regulatory capacity year-round while primarily regulating specific targets to match seasonal demands, allowing rapid transcriptional changes without the energetic cost of transcriptional overhaul (Ilker & Hinczewski, 2024). Similar post-transcriptional robustness is documented in invertebrates and insects that survive extreme thermal fluctuations and anoxia (Biggar et al., 2012; Lyons et al., 2016). Likewise, miRNAs serve a critical role in plant responses to elevated temperatures and drought (Samad et al., 2017). Importantly, this flexibility in regulation allows the coral to orchestrate complex physiological transitions over the course of the year, modulating the transcriptional landscape to fulfill winter metabolic quiescence, energetic demands of gametogenesis, and homeostasis.

The regulatory capacity of miRNA supports homeostasis and repair at the thermal extremes of the annual cycle (February and August). miRNA regulation of mRNA was associated with suppression of Wnt signaling and structural morphogenesis (Fig. 5A, B). Wnt signaling is a master regulator of body patterning and axis formation in cnidarians (Uveira et al., 2025; Vetrova & Kremnyov, 2025), and its selective suppression suggests that *A. poculata* uses a miRNA-mediated strategy to halt resource-intensive structural organization during periods of low and high temperatures. This seasonal suppression in August also likely mitigates cellular damage from thermal and UV stress (Courtial et al., 2017). In February, miRNAs were specifically associated with the repression of mRNAs involved in cell cycle progression and protein turnover (Fig. 5C, D). By suppressing cell division and ubiquitin-mediated protein degradation, *A. poculata* appears to be stabilizing its existing protein pool rather than incurring the energetic cost of turnover (Glickman & Ciechanover, 2002; Peth et al., 2013), a hallmark of metabolic depression in other dormant organisms (Grabek et al., 2015; Ramnanan et al., 2009). Similar miRNA-driven enforcement of quiescent states has been described across diverse taxa, from mammalian hibernation to invertebrate diapause, where specific miRNAs act as checkpoints controlling apoptosis, cell cycling, and developmental timing (Arfat et al., 2018; Biggar et al., 2012; Hadj-Moussa & Storey, 2020; Reynolds, 2019; Zhao et al., 2015).

Certain mRNA-miRNA interactions showed stable negative correlations across all three time points, including those related to calcium homeostasis (Fig. 5I, J). Calcium is a universal second messenger controlling a wide range of cellular functions (Clapham, 2007; Fedrizzi et al., 2008), and disrupted calcium signaling can trigger cellular dysfunction and death (Ermak & Davies, 2002; Brini et al., 2014). By maintaining constant repression of genes related to calcium, miRNAs may be protecting cells from ‘leaky’ or spurious calcium spikes that could inadvertently trigger premature stress responses. This stabilizing role is particularly critical in scleractinian corals, where calcium dynamics are central to calcification and the bleaching signaling cascades (Helgoe et al., 2024). These constitutive interactions represent a robust regulatory infrastructure that ensures cellular homeostasis is maintained even as other network components are dynamically rewired to meet seasonal demands in *A. poculata*.

In June specifically, miRNA regulation of mRNAs was associated with the repression of GABA signaling and microvillus assembly (Fig. 5G, H). While GABA is well known for its role in the settlement and metamorphosis of marine invertebrates (Meyer et al., 2011; Ishii et al., 2022; Guo et al., 2025), it may also act as a neuro-endocrine signal coordinating reproductive timing, as seen in other taxa (Cao & Ling, 2025; Kusuma & Hariani, 2017); therefore, its repression in June may serve as the final stages of gamete maturation or a spawning regulatory cue in *A. poculata*. In parallel, the repression of microvillus assembly suggests either a shift in nutritional strategy (Lindsay et al., 2025; Trumbauer et al., 2021) or a gametogenesis endpoint (Goldberg, 2002; Raz-Bahat et al., 2017; Valente et al., 2024) as light and temperature increase.

A notable and recurring finding across multiple clusters (Clusters 2, 3, and 4; Fig. 5) was the functional enrichment of miRNA-mediated gene silencing and miRNA metabolic processes among targets negatively correlated with miRNAs. This meta-regulatory pattern, where miRNAs appear to target components of the very machinery that produces and deploys them, including Dicer and Argonaure homologs, has been documented in model systems (Tokumaru et al., 2008; Martello et al., 2010; Finnegan & Pasquinelli, 2013; Liu et al., 2021; Suster & Feng, 2021), but is described here for the first time in a coral. This suggests that *A. poculata* employs miRNA to fine-tune its own regulatory miRNA machinery, indicating that transcriptional responses to environmental variation remain within cellular homeostatic limits. This finding positions miRNAs as central coordinators of transcriptional plasticity, capable of modulating the broader regulatory infrastructure itself.

#### MicroRNA networks reorganize proactively to buffer against thermal stress without phenotypic changes

The +3°C treatment produced negligible physiological effects and only minimal differential gene expression relative to ambient conditions at either June or August time points. This apparent maintenance of homeostasis under moderate warming aligns with previous observations of thermal resilience in this species (Allen-Waller et al., 2025; Harman et al., 2025; Wuitchik et al., 2021). Notably, this homeostasis was not maintained by large-scale *de novo* transcription of stress genes but instead by a flexible re-wiring of the mRNA-miRNA regulatory network. Under heat conditions in both June and August, the miRNA regulatory network reorganized such that targets associated with development, signaling, and protein turnover, demonstrated repression patterns. These processes are not typically classified as canonical heat stress responses, but may instead reflect proactive cellular reconstruction to offset potential thermal stress before physiological signs of stress emerge. By reconfiguring regulatory networks, miRNAs can confer cellular robustness, stabilizing physiology without necessitating a change in observable phenotype (Cassidy et al., 2013; Leung & Sharp, 2007, 2010; Siciliano et al., 2013).

## Conclusions

This study provides the first characterization of the miRNA repertoire in *Astrangia poculata*, and demonstrates that seasonal temperature variation, rather than moderate chronic warming, is the dominant driver of physiological and molecular landscape. By establishing this seasonal transcriptomic baseline, we demonstrate that *A. poculata*’s response to seasonal cycles is far more expansive than its response to moderate thermal challenges, suggesting that the inherent plasticity required for seasonal survival may serve as a critical buffer against climate change. Our findings demonstrate that miRNAs are both a broad regulator of plasticity, but also fine tune specific and seasonally distinct gene expression. These results indicate quiescence is a coordinated molecular state rather than passive metabolic inactivity, an important distinction for our understanding of sessile marine invertebrates. Furthermore, this miRNA-mediated framework provides stable yet flexible infrastructure that allows *A. poculata* to absorb a +3°C warming challenge without necessitating a discrete stress response. Together, these findings illustrate coordinated seasonal transcriptomic shifts: winter emphasizes damage prevention and cellular integrity, early summer prioritizes reproductive and growth investment, and late summer underscores metabolic flexibility, coupled with heightened immune activity. These shifts highlight how northern populations of *A. poculata* balances quiescence, reproduction, growth and defense to survive in seasonally variable environments. However, whether this regulatory plasticity can be sustained as climate change pushes thermal maxima beyond historical thresholds remains an open and pressing question.

## Acknowledgments

JA was supported by the Graduate Research Fellowship from the National Science Foundation and the Global Marine Initiative Student Research Award from The Nature Conservancy. CG was supported by the University of Rhode Island Coastal Fellowship program. *Astrangia poculata* were collected under Rhode Island Department of Environmental Management collector’s permit #970. The authors extend appreciation to the Temperate Coral Research Conferences hosted by Roger Williams University, Tufts University, and Southern Connecticut State University for fostering creative conversations and collaborations. We thank Anya Hanson and the URI dive program, along with the following individuals for assistance in coral collections: Cassie Raker, Taylor Lindsay, Jack Girard, Matias Gómex-Corrales, Danielle Becker, Alex Moen, and Anya Hanson. Edward Baker provided facility support at the Graduate School of Oceanography. We thank Alice Ball, Katie Barott, Danielle Becker, Ben Glass, Megan Guidry, Sam Gurr, Ariana Huffmyer, Hannah Reich, Emma Strand, Kevin Wong, and Amy Zyck for experimental support. We extend much appreciation to Zoe Dellaert and Taylor Lindsay for analytic support, as well as Zoe Dellaert and Federica Scucchia for feedback on manuscript drafts.

## Data availability

Data and code to reproduce the analyses presented here are available at https://github.com/JillAshey/Astrangia_repo/. RNA sequences and small RNA sequences can be found on NCBI under BioProjects PRJNA1231118 and PRJNA1231129, respectively.

