## Supplemental Figures for "A miRNA-mediated gene regulatory network supports seasonal plasticity in the temperate coral *Astrangia poculata*"

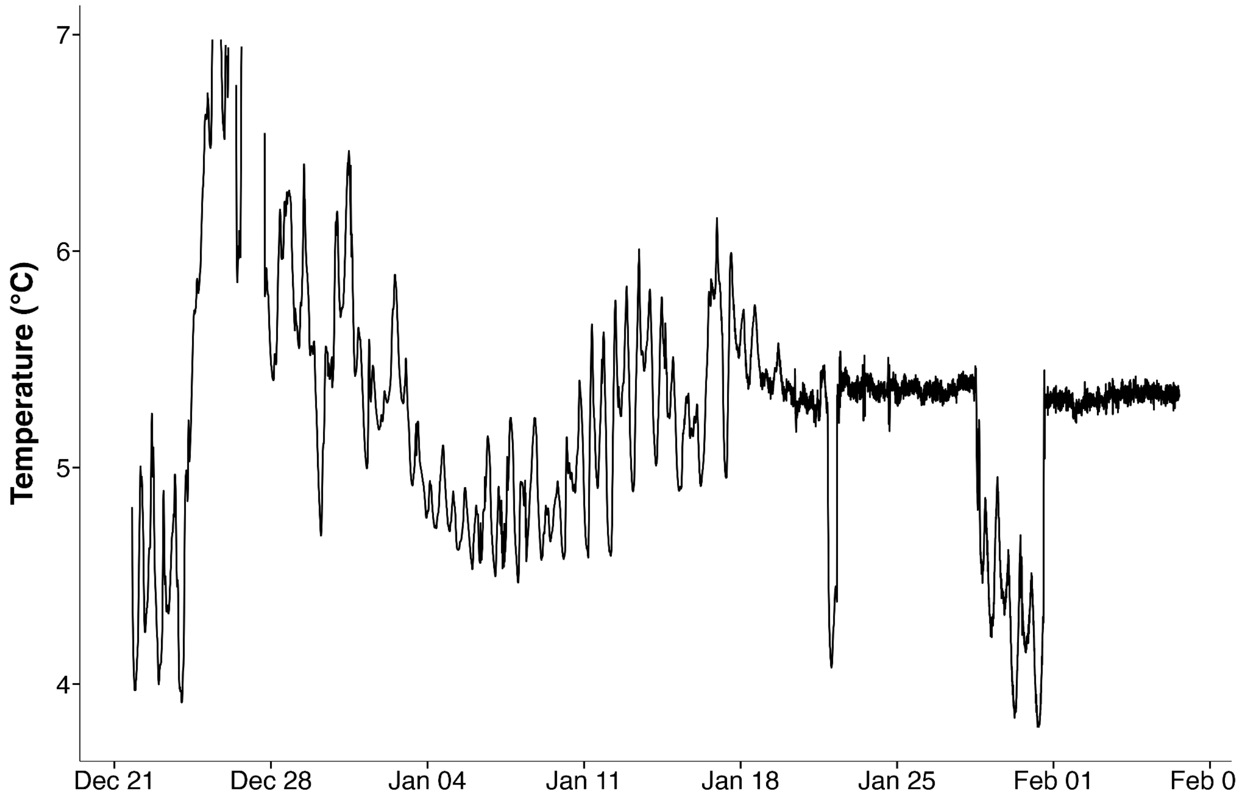


**Figure S1.** Temperature (°C) during the acclimation period (December 2020 - January 2021).

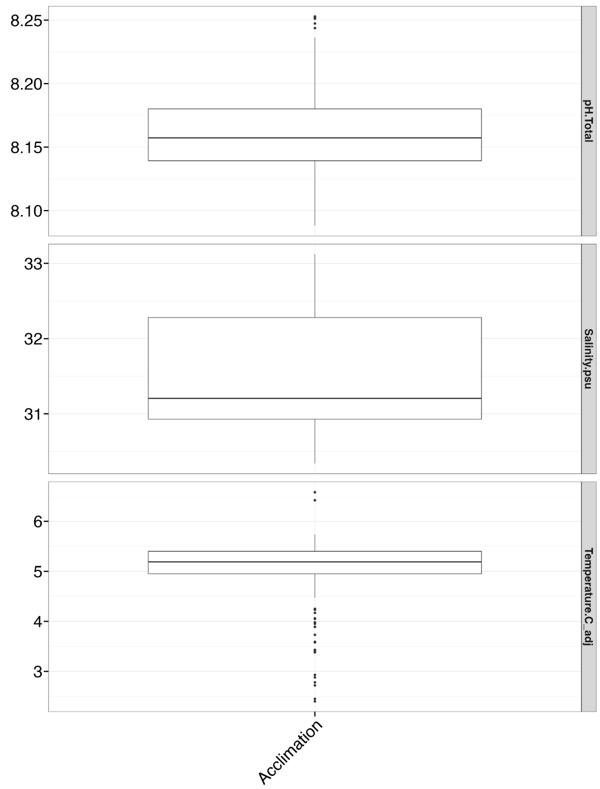


**Figure S2.** Discrete measurements over the course of the acclimation period (December 2020 - January 2021). The top panel displays pH (total), the middle panel displays salinity (psu), and the bottom panel displays temperature (°C).

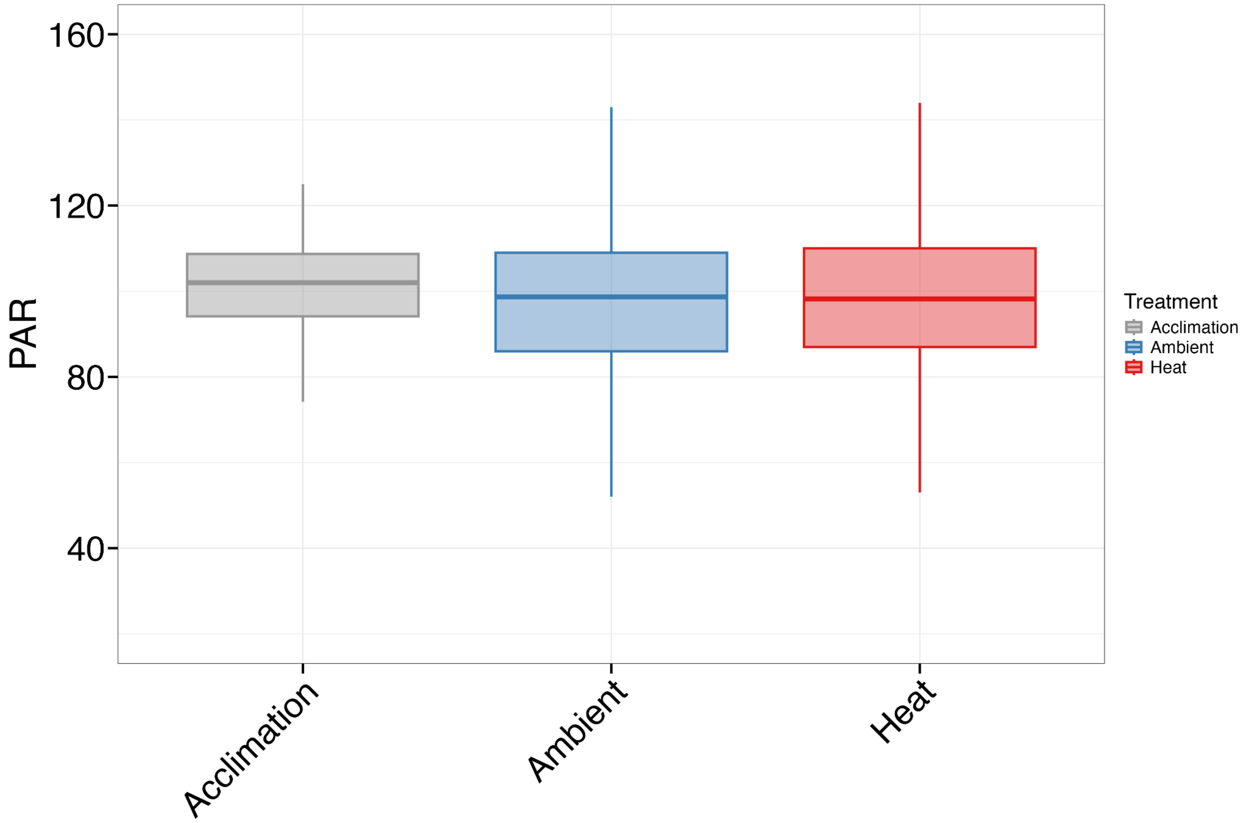


**Figure S3.** Light data from acclimation, ambient and heat treatments. Light reported in PAR measured with an Underwater Quantum Meter (Model MQ-510: Apogee, µmol $m^{-2}$ $s^{-1}$) as described in the methods.

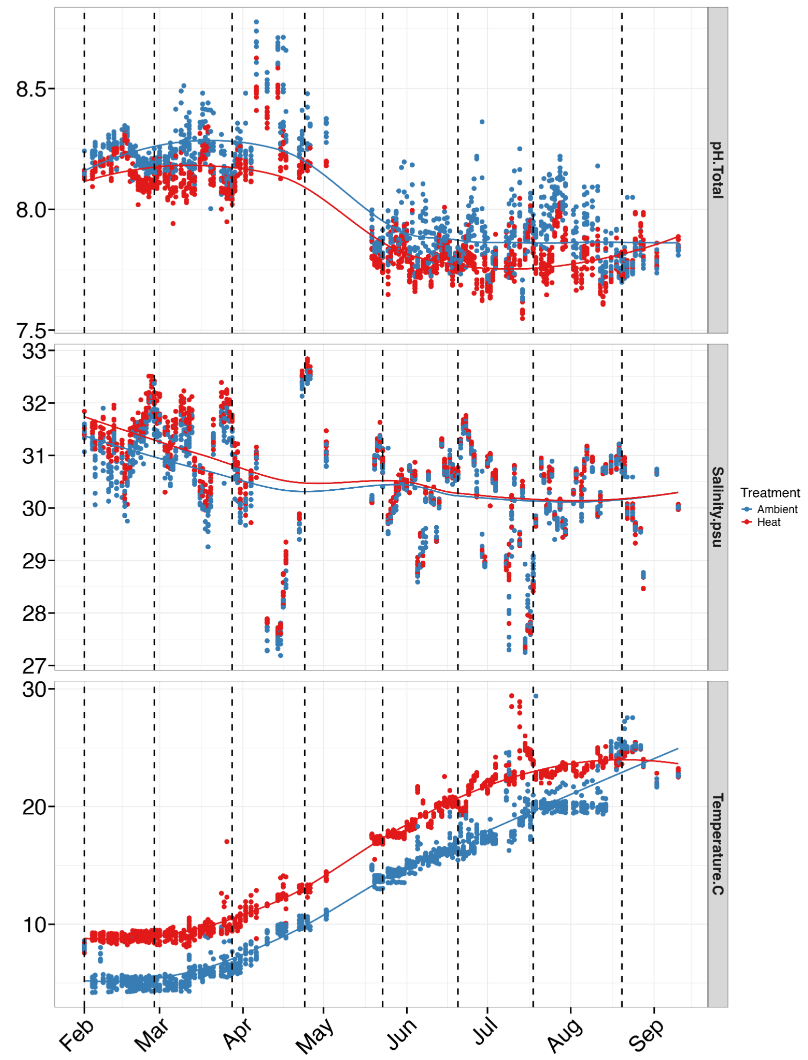


**Figure S4.** Discrete measurements over the course of the experimental period (February 2021 - August 2021). The top panel displays pH (total scale), the middle panel displays salinity (psu), and the bottom panel displays temperature (°C). Dotted vertical lines indicate sampling time points.
